# Taming the Genetic Fire: Transposable Element diversity across thermal environments in polychaetes

**DOI:** 10.64898/2026.02.25.703748

**Authors:** Laure Lamothe, Stéphane Hourdez, Thierry Robert, Éric Bonnivard

## Abstract

Genetic variation plays a central role in enabling organisms to adapt to ever-changing environments. Transposable elements (TEs) are key drivers of genetic variation and adaptation, partly due to their ability to respond to environmental changes, such as thermal variability, through transcriptional activation, potentially leading to insertion events. The new copies will eventually accumulate mutations, increasing the TE diversity in the genome. In this study, we investigated how the TE diversity varies across environments, contrasted by their average temperature and their thermal variability profile, using polychaete annelids as a model system. These primarily benthic organisms occupy a wide range of habitats, from polar waters to hydrothermal vents and temperate shores. TE diversity varied substantially among polychaete species, with significantly lower diversity observed in species inhabiting unstable habitats, such as those associated with hydrothermal vents. This link between TE diversity and environment was statistically consistent across the different TE orders, except for DIRS-like elements in Errantia polychaetes, that display a surprisingly high diversity. Our results suggest that TE diversity may be selected to balance the level of TE activation, linked to thermal variability, to maintain a sustainable mutation rate at the whole genome level. In unstable environments, high TE diversity may not be sustainable due to the accumulation of deleterious mutations, caused by a higher rate of stress-induced transposition compared to other habitats. These findings highlight the influence of environmental conditions on the long-term dynamics governing TE-host interactions and underscore the role of TEs in evolution.

## INTRODUCTION

Phenotypic plasticity and/or genetic variation are required to face new environmental conditions (King et al., 2018; Stamp & Hadfield, 2020; Valladares et al., 2014). As ubiquitous mobile genetic elements, transposable elements (TEs) are a prominent source of genetic variation, particularly since transposition can induce a wide range of mutations. They allow the emergence of new promoters, trigger exon shuffling and chromosomal rearrangements, and reshape regulatory systems of their hosts (Bourque et al., 2018; Feschotte, 2008; Gebrie, 2023; Schrader & Schmitz, 2019). Formerly considered “junk DNA” or genetic parasites, TEs are now recognized as major contributors to phenotypic evolution. However, the host-TE relationship is complex and cannot be reduced to simple parasitism or mutualism (Chakrabarty et al., 2023; Cosby et al., 2019). Although some TE-induced mutations may confer a selective advantage to the host, most are neutral or deleterious, so regulatory systems repressing transposition are essential to limit the mutation load at the whole genome level (Ayarpadikannan & Kim, 2014). A typical feature of long-term host-TE interactions is therefore coevolution toward a balance between transposition and repression or silencing (Yoth et al., 2022). Only TEs capable of temporarily escaping repression mechanisms and being transcribed may maintain active copies in genomes, in populations and eventually at the species level (Blumenstiel, 2019; Redd et al., 2023; Sammarco et al., 2022). The other TEs progressively lose functionality, decay, and ultimately disappear from the genomes. A well-documented way for TEs to escape repression is through stress-induced activation during a biotic or abiotic stress (Capy et al., 2000 Chadha & Sharma, 2014; Grandbastien et al., 2005), the colonization of new environments by the host (Kalendar et al., 2000; Vieira et al., 1999), or a change in host silencing mechanisms (Guerreiro, 2012; Pappalardo et al., 2021). This stress responsiveness promotes genetic novelty, which may even lead to the radiation of new taxa (Belyayev, 2014), and plays a significant role in adaptive responses (Casacuberta & González, 2013; Schrader & Schmitz, 2019; Yuan et al., 2022). Among abiotic factors, temperature is a known contributor to TE activation. However, most studies have focused on terrestrial model species (Bogaerts-Márquez et al., 2021; Cayuela et al., 2022; Guerreiro, 2012; Jardim et al., 2015; Wos et al., 2021). Given the strong influence of temperature in shaping the physiology and biogeography of marine species (Helaouët & Beaugrand, 2009) and considering the ongoing disruptions of ocean temperatures caused by anthropogenic climate change (Garcia-Soto et al., 2021), understanding how marine organisms adapt to their thermal environment is essential. Such knowledge would not only clarify how temperature has contributed to shape their evolutionary history but also improve predictions of how marine life will respond to global environmental changes. In the marine environment, the case of notothenioid fish, which arose through adaptive radiation during Southern Ocean cooling, has caught the interest of the scientific community. This radiation has been associated with bursts of TEs (Auvinet et al., 2018; Bista et al., 2023; L. Chen et al., 2019; Z. Chen et al., 2008). Besides fish, studies have mainly focused on differential expression of TEs under heat stress. Some have revealed thermal stress-induced TE activation in shrimp (de la Vega et al., 2007), corals (Desalvo et al., 2008; Rose et al., 2016), and bivalves (Lesser et al., 2019). Nevertheless, the influence of temperature on TEs in the marine environment remains largely understudied, especially with respect to TE diversity.

TEs are highly diverse elements, diverging first in their mode of transposition, which likely reflects distinct evolutionary origins (Wells & Feschotte, 2020). Retrotransposons are reverse transcribed from an RNA intermediate before insertion, whereas DNA transposons use a DNA intermediate to transpose. Within each of these TE classes, the high diversity of structures, functional domains, and transposition mechanisms differentiates TE orders (reviewed in Wicker et al. 2007). These orders also vary greatly in their distribution, diversity, abundance, and impact on host genomes. Four orders of retrotransposons are well defined: (i) LTR-retrotransposons, (ii) DIRS-like retrotransposons (DIRS and Ngaro), (iii) Long Interspersed Nuclear Elements (LINEs), and (iv) Penelope-like elements. Two orders of DNA transposons are described: (i) TIR/Crypton elements and (ii) Maverick/Helitron. Within each order, TEs exhibit high structural and sequence diversity, which defines TE superfamilies — 60 of which have been characterized (https://www.girinst.org/repbase/) and are considered in this paper. At a finer scale, superfamilies are subdivided into families, each grouping all copies of the same element. There is a two-way link between the diversity of TEs and their activity. Transposition increases the number of copies, which can then diverge and give rise to new families (Wells & Feschotte, 2020). Conversely, a genome with highly diverse TEs, particularly in their promoters and regulatory sequences, might present a higher chance that some TEs escape repression and transpose, which can result in higher overall TE activity. Using the number of families as an estimate of TE diversity (Thomas-Bulle et al., 2018), we aimed to explore, in marine animals, the relationships between TE diversity at the family level (TDF) and host thermal environment. Polychaeta, a class of worms living mainly in marine benthic habitats from intertidal to abyssal depths, is a suitable model because its species are found in highly contrasted habitats, ranging from polar regions to deep-sea hydrothermal vents and coastal environments. An additional advantage is that the two main subclasses, Errantia (usually free-living) and Sedentaria (sedentary or tube-dwelling), are well represented across these diverse habitats (Struck et al., 2011). Assessing TDF across many species from the same taxonomic group requires large amounts of OMIC data. However, complete genome sequences are still lacking for polychaetes outside of model species. Here, we took advantage of extensive transcriptomic resources available in public databases to conduct a large comparative study of 50 polychaete species. We developed DetecTE2, an improved version of the first tool able to estimate TDF from transcriptomes (Filée et al., 2021) and used it to investigate the links between TDF and host thermal environment while accounting for the phylogenetic structure of the dataset.

## METHODS

### Transcriptomic data selection and metadata

We selected 50 species from around the globe (Figure 1), which represent the taxonomic diversity of marine polychaetes (Figure 2). The study was restricted to non-cosmopolitan species from contrasting environments. Each species should unambiguously be assigned to one of five thermal environments (Supplementary Material 1), defined by temperature characteristics in their natural habitat: (1) Stable-cold: average annual temperatures below 5 °C with low variation (less than 8 °C annually), corresponding to deep-sea (Emery, 2001) and polar regions (https://atlas.climate.copernicus.eu/atlas), (2) Stable-hot: average annual temperatures between 25 °C and 35 °C with low variation (less than 8 °C annually), characteristic of coasts at low latitudes (https://atlas.climate.copernicus.eu/atlas), (3) Cyclic-intermediate: average annual temperatures between 10 °C and 15 °C with seasonal variations (more than 8 °C annually), found in coastal regions of temperate zones (https://atlas.climate.copernicus.eu/atlas), (4) Unstable-hot: average temperatures between 25 °C and 35 °C with stochastic variations (more than 8 °C), characteristic of hydrothermal vents at fluid outlets (Fustec et al., 1987), (5) Unstable-intermediate: average temperatures between 10 °C and 15 °C with stochastic variations (more than 8 °C), found in diffuse hydrothermal vents or areas distant from hydrothermal fluid outlets (Fustec et al., 1987). The transcriptome of one individual by species was studied. The analyzed transcriptomes can be divided into two distinct sets. Twenty-five of them had already been studied with respect to the diversity of transposable elements, particularly LTR-retrotransposons, in polychaete annelids (Filée et al., 2021). Twenty-five additional transcriptomes were selected through data mining in NCBI and added to this first dataset (Supplementary Material 1). Transcriptomes were selected with at least 35 million paired-end Illumina reads to ensure reliable detection of TEs (Filée et al., 2021). Contaminated data, as indicated by the read analyses provided on NCBI website, were excluded. We also verified that the sequenced tissues were sampled from natural environments without any specific experimental treatments prior to RNA extraction.

**Figure 1:**
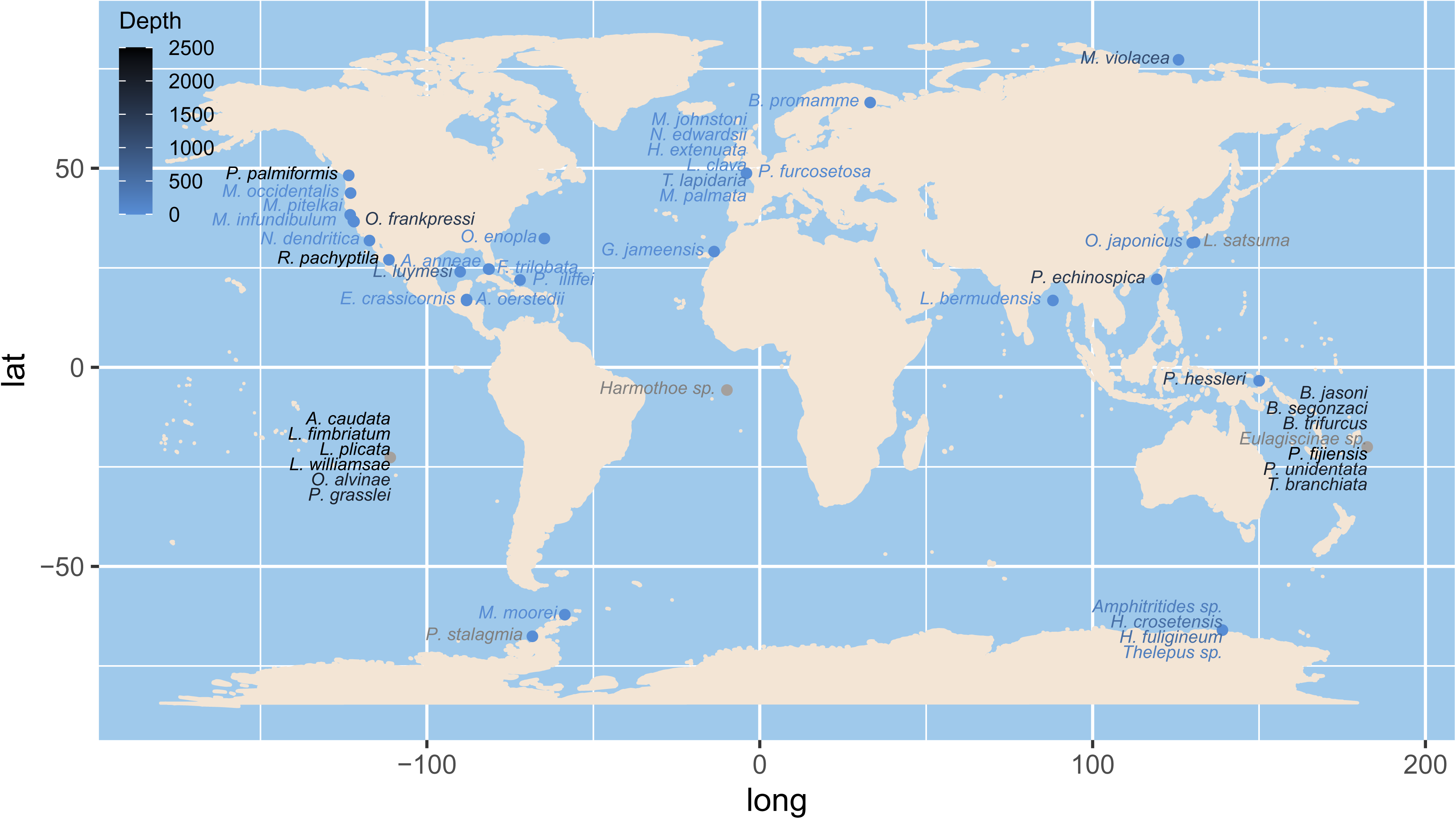
Map of sample collection sites. Species names are colored by sampling depth. Grey point indicates imprecise location due to missing coordinates.

**Figure 2:**
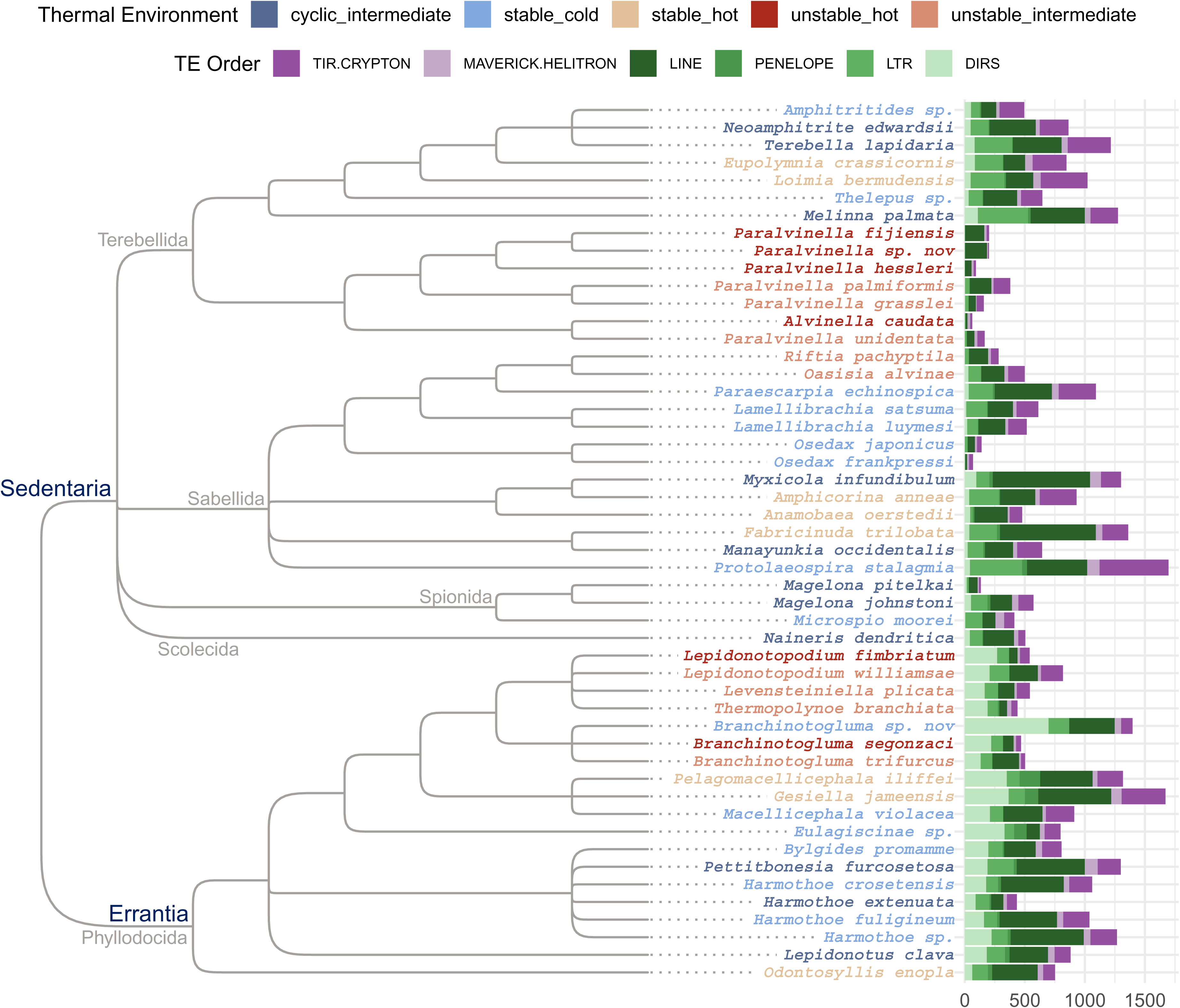
TE order-specific TDF by species. Left : Cladogram of the phylogeny of the 50 polychaete species, based on Sun et al. 2018, Stiller et al. 2020, Tilic et al. 2020, Gonzalez et al. 2023, and Brun et al. 2024. Right : Number of active TE families detected from species transcriptomes.

### Assembly of transcriptomes

To remove low-quality bases and reads, FastP (https://github.com/OpenGene/fastp) was run with default settings on each set of reads. Read quality was then assessed using FastQC with default parameters (https://github.com/s-andrews/FastQC). The cleaned data were subsequently down sampled to 40 million high-quality reads (±10%), except for *Branchinotogluma segonzaci* (33M) and *Branchinotogluma* sp. (33M). Transcriptome assemblies were generated independently using two assembly tools to maximize subsequent TE detection: Trinity (https://github.com/trinityrnaseq/trinityrnaseq/wiki) and RNAspades (https://github.com/ablab/spades). Both were run with 20 threads, 150 GB of memory, and default parameters.

### Detection and Annotation of Transposable Elements

Our first tool allowing the annotation of TEs from transcriptome data without a reference genome or mobilome (Filée et al., 2021), hereafter referred to as DetecTE1, was optimized for this study and released as DetecTE2 (https://github.com/laurelamothe/DetecTE2). DetecTE2 extracts the actual TE sequence instead of the whole transcript and merges TE sequences based on a chosen level of similarity. DetecTE2 can also handle multiple transcriptome inputs, which are analyzed independently up to and including the fifth step (see below for the description of the DetecTE2 pipeline). DetecTE2 requires as input a protein database of TEs in FASTA format. In this study, the database corresponds to the LAC28 database (Filée et al., 2021) enriched with 517 annelid TE sequences (Supplementary Material 2) detected in the transcriptomes of *Alvinella caudata, Neoamphitrite edwardsii, Amphitritides sp., Harmothoe sp., Melinna palmata, Paralvinella fijiensis, Paralvinella palmiformis, Paralvinella unidentata, Paralvinella hessleri*, and *Paralvinella grasslei* by tblastn against the curated database (e-value < 1e-21, pident > 25%, length > 50 aa, gapopen < 15). These sequences were then validated with CENSOR on Repbase, to check the consistency of the annotation and remove chimeric sequences, before being added to the database. DetecTE2 proceeds as follows (Supplementary Material 3). (1) Transcripts shorter than 500 bp are removed. (2) Transcripts carrying TEs are identified using crossmatch *via* BLAST (https://ftp.ncbi.nlm.nih.gov/blast/executables/blast+/2.14.0/) against the protein database. Only transcripts with hits belonging to the same superfamily in both blastx (e-value < 1e-21) and tblastn (e-value < 1e-21, pident > 25%, length > 50 aa, gapopen < 15) are selected. (3) TE extraction: The positions of TEs in each transcript are extracted from the tblastn output. As BLAST usually produces redundant hits, overlapping hits on the same transcript are concatenated. Additionally, hits belonging to the same TE superfamily and separated by fewer than 250 amino acids are concatenated. (4) TE annotation: For each TE sequence, if at least eight of the ten best tblastn hits belong to the same superfamily, the TE is annotated accordingly. Otherwise, it is annotated as “Unknown”. (5) TE sequences shorter than 500 bp are removed. (6) TE families identification: All extracted TE sequences from the input transcriptomes are compared against each other using blastn (e-value < 1e-21, pident > 90%, length > 100 bp). TE sequences are then grouped into families, defined as connected components based on the blastn results. A family therefore includes all sequences with at least 90% identity over 100 bp pairwise. If at least 80% of the sequences belong to the same superfamily, the family is annotated accordingly. Otherwise, it is annotated as “Unknown”. (7) The number of families per superfamily is counted and recorded in the output table. We used the number of TE families, the lowest level of TE classification, as a measure of TE diversity, as it allowed a better resolution for the subsequent analyses of the differences in TE diversity between species.

Taking advantage of the few publicly available polychaete genomes, we used RepeatModeler to estimate the TDF from the genomes of *Lepidonotus clava* (GenBank accession: GCA_936440205.1), *Osedax frankpressi* (GCA_964035775.1), *Paraescarpia echinospica* (GCA_020002185.1), *Terebella lapidaria* (GCA_949152475.1), and *Magelona johnstoni* (GCA_963942565.1) and compared it with the TDF estimated for the same species from transcriptomes with DetecTE2. Default parameters were used, and the TDF was defined as the sum of TE families annotated as DNA, LINE, or LTR by RepeatModeler.

### Statistical Analyses

All statistical analyses were performed using R (version 4.3.2). The TDF estimated by DetecTE2 was compared to the estimates of DetecTE1 and RepeatModeler on whole genomes using Spearman correlation (stats package, version 3.6.2). To describe TDF variation according to thermal environments and phylogenetic groups, a principal component analysis (PCA) was conducted (FactoMineR package, version 2.9). To further assess the effect of the thermal environment on TDF, non-parametric Kruskal&Wallis tests (stats package, version 3.6.2) were conducted to cope with the low sample size in some of the environments. Pairwise rank comparisons were carried out using Nemenyi tests (FSA package, version 0.9.6). In addition, for each TE order, a linear regression of the TDF according to the thermal environment was also fitted using a phylogenetic generalized least squares (pGLS) model (nlme package, version 3.1-168), to account for the non-independency of the samples due to phylogenetic relationships between species. The pGLS used an evolution model derived from Brownian motion (ape package, version 5.8-1) with, as input, the covariance matrix of error terms derived from the cladogram shown on Figure 2. For each analysis, in order to check the convergence of Pagel’s λ estimator, repetitions of the iterative estimation procedure were done using 100 initial values from zero to one. Convergence of λ estimations was systematically observed. In order to test the significance of the phylogenetic signal, each pGLS model was compared to the corresponding GLS model which ignores the phylogenetic relationship between species (Pagel’s λ = 0). The R² of the models based on the log-likelihood estimators of sum of squares were computed using the rr2 package (version 1.1.1). Tukey HSD post hoc test (emmeans package, version 1.11.2-8) were performed for pairwise mean comparisons. A Kendall’s trend test (irr package, version 0.84.1) was performed to evaluate the consistency of the ranks of TDF means by thermal environments across every TE order. The differences in DIRS-like TDF between Errantia and Sedentaria species, as well as the influence of the thermal environment on TDF for Errantia and Sedentaria species considered separately were assessed with Kruskal–Wallis tests (stats package, version 3.6.2). No GLS or pGLS analysis was performed on the Errantia and Sedentaria subsamples respectively, due to insufficient numbers of species. Pairwise comparisons were performed using Nemenyi tests (FSA package, version 0.9.6). All Kruskal–Wallis tests were carried out after checking the equality of rank variances using a Fligner test (stats package, version 3.6.2). In case of rank variance inequality, a log transformation was performed for the Fligner and Kruskal&Wallis tests (indicated by a star in Figure 4).

## RESULTS

TDF estimated by DetecTE2 and DetecTE1 (from Filée et al., 2021) were compared. The estimates were reduced by 38% on average in DetecTE2, but the TDF estimated by the two tools showed a strong correlation (Spearman coefficient = 0.927, Supplementary Material 4). Considering the comparison of TDF estimates from whole genomes and transcriptomes, the TDF was on average 50% higher when assessed from whole genomes (Supplementary Material 5) but also highly correlated with TDF in transcriptomes (Spearman coefficient = 0.9).

### Transposable Element Diversity in Polychaetes

TDF was first estimated for each superfamily (Supplementary Material 6) and then grouped by TE order (Figure 2). Total TDF in polychaetes had a mean value of 716 families and was highly variable among species, with a coefficient of variation of 61%. It ranged from only 63 families in *A*. *caudata*, a thermophilic hydrothermal Terebellida (Desbruyères et al., 2006), to 1697 in *Protolaeospira stalagmia*, an Antarctic Sabellida (Nieva et al., 2021). Furthermore, TDF in polychaete species varied substantially among the six TE orders (Table 1). The mean TDF of Penelope-like, Helitron/Maverick, DIRS-like, LTR-retrotransposons, and TIR/Crypton DNA transposons was 18, 40, 107, 124, and 157 families, respectively. Finally, LINEs were by far the most diverse TE order, with an average TDF of 271 families and the highest TDF in 70% of the species. The distribution of families also differed according to the TE orders. Penelope-like and DIRS-like elements showed a patchy distribution, being absent in 16% and 14% of species respectively, even though the average TDF of DIRS-like elements is six times higher than that of Penelope-like elements. In contrast, Helitron/Maverick, TIR/Crypton elements, and LINEs were ubiquitous despite their drastically different average TDF. Interestingly, a clear differentiation between Errantia and Sedentaria species for TE diversity is observed, as Errantia species displayed significantly higher TDF (Kruskal–Wallis H = 5.23 ; p-value = 0.022), with a median of 892 families per species compared to 609 families in Sedentaria. DIRS-like diversity mainly contributed to this difference (Figure 2 and Figure 3). These elements displayed a high TDF in Errantia (232 by species on average), whereas their number of families remained low in most Sedentaria species (31 by species on average ; Kruskal–Wallis H = 33.34 ; p-value = 7e-9). In particular, the Sedentaria included seven species with no DIRS-like elements detected, as well as four species with five families or fewer. The diversity of the Penelope-like elements, represented by five families or fewer in half of the dataset, also appeared to be higher in Errantia, with a mean of 14 families compared to 5 in Sedentaria, but the comparison of medians was not significant (Kruskal–Wallis H = 1.07 ; p-value = 0.302).

**Figure 3:**
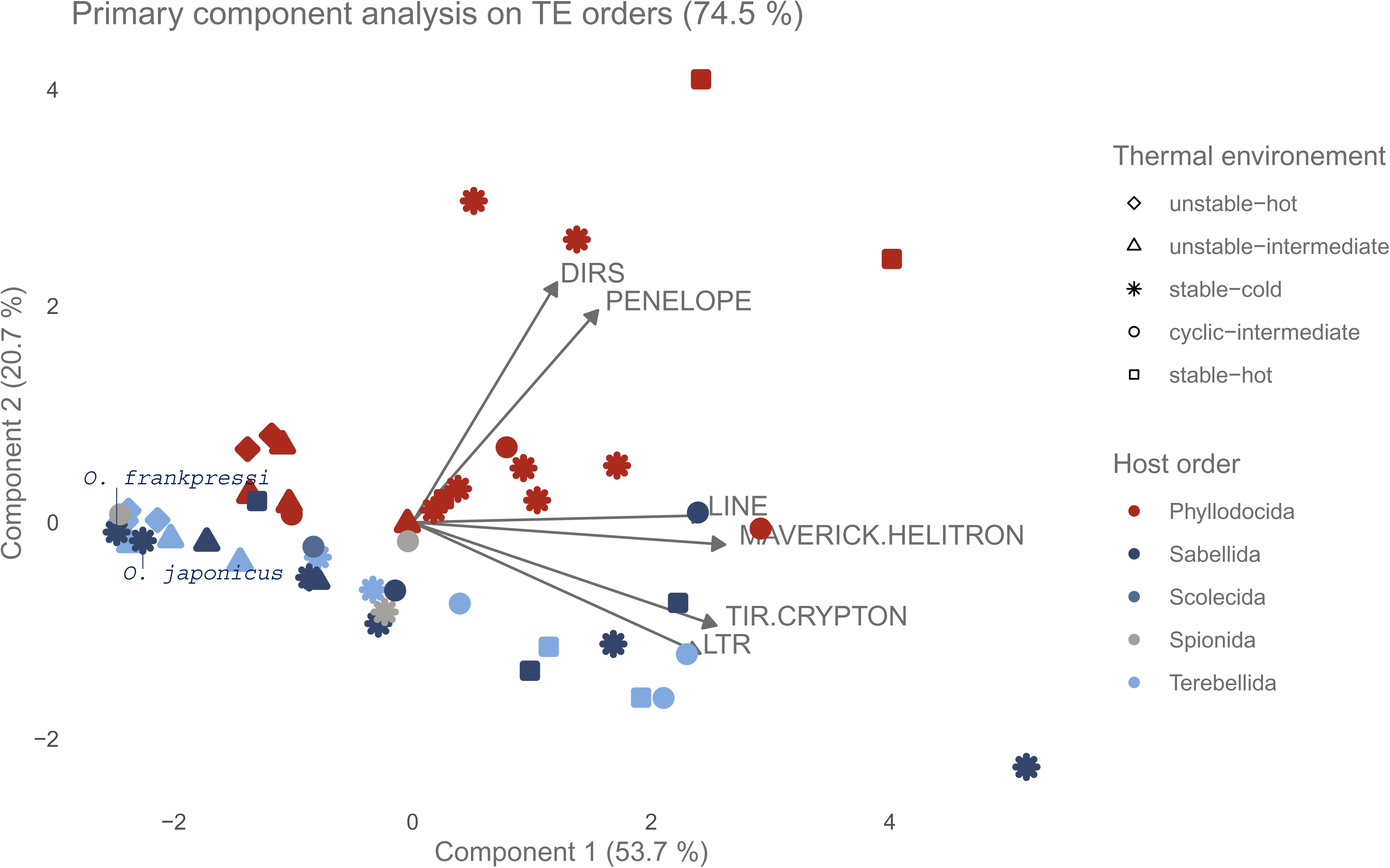
Primary Component Analysis of the TE order-specific TDF by species. The arrows show the projection of the six response variables (TDF by order) on the first and second components. The contributions of the first and second components to the global variance are written on the x and y axis labels, respectively. DIRS-like, Penelope-like, LINEs, Maverick/Helitron, TIR/Crypton, and LTR retrotransposons order have a correlation of 0.403, 0.518, 0.816, 0.872, 0.849 and 0.804 with the first component respectively and of 0.739, 0.653, 0.021, −0.068, −0.318, −0.404 with the second component respectively.

### Patterns of total TE Diversity According to the Thermal Environment

The pattern of TDF across the different thermal environments and host taxonomy is illustrated in Figure 3. Specifically, species from thermally unstable environments, both hot and intermediate, were clearly distinguished on the first dimension of the PCA from species from the stable-hot environment. LINEs, Helitron/Maverick, LTR-retrotransposons, and TIR/Crypton DNA transposons contributed mainly to this difference, all being more diverse in species from the stable-hot environment. On the contrary, species from cyclic-intermediate and stable-cold environments were highly scattered on both dimensions of the PCA. This pattern was statistically supported by a significant relationship between the global TDF and the thermal environment, mainly due to significant differences between species from unstable environments on one side, and stable or cyclic environments on the other side (Figure 4 and Supplementary material 7). The stable-cold environment displayed the highest variation in TDF. In particular, Sabellida species in stable-cold environments appeared widely dispersed (Figure 3). Indeed, for annelids from the stable-cold environment, Sabellida included both the species with the highest TDF (1697 families in *P. stalagmia*, Supplementary material 6) and two of the five species with the lowest TDF (69 families in *O*. *frankpressi* and 140 in *Osedax japonicas*, Supplementary material 6). These two bathypelagic bone-eating worms appeared strongly differentiated from other species living in the stable-cold environment, clustering with hydrothermal Terebellida species (Figure 3). Similarly, species living in cyclic-intermediate environments showed strong differences in their TDF (Figure 3), although the small number of species in each phylogenetic group did not allow identification of taxonomy-related characteristics.

**Figure 4:**
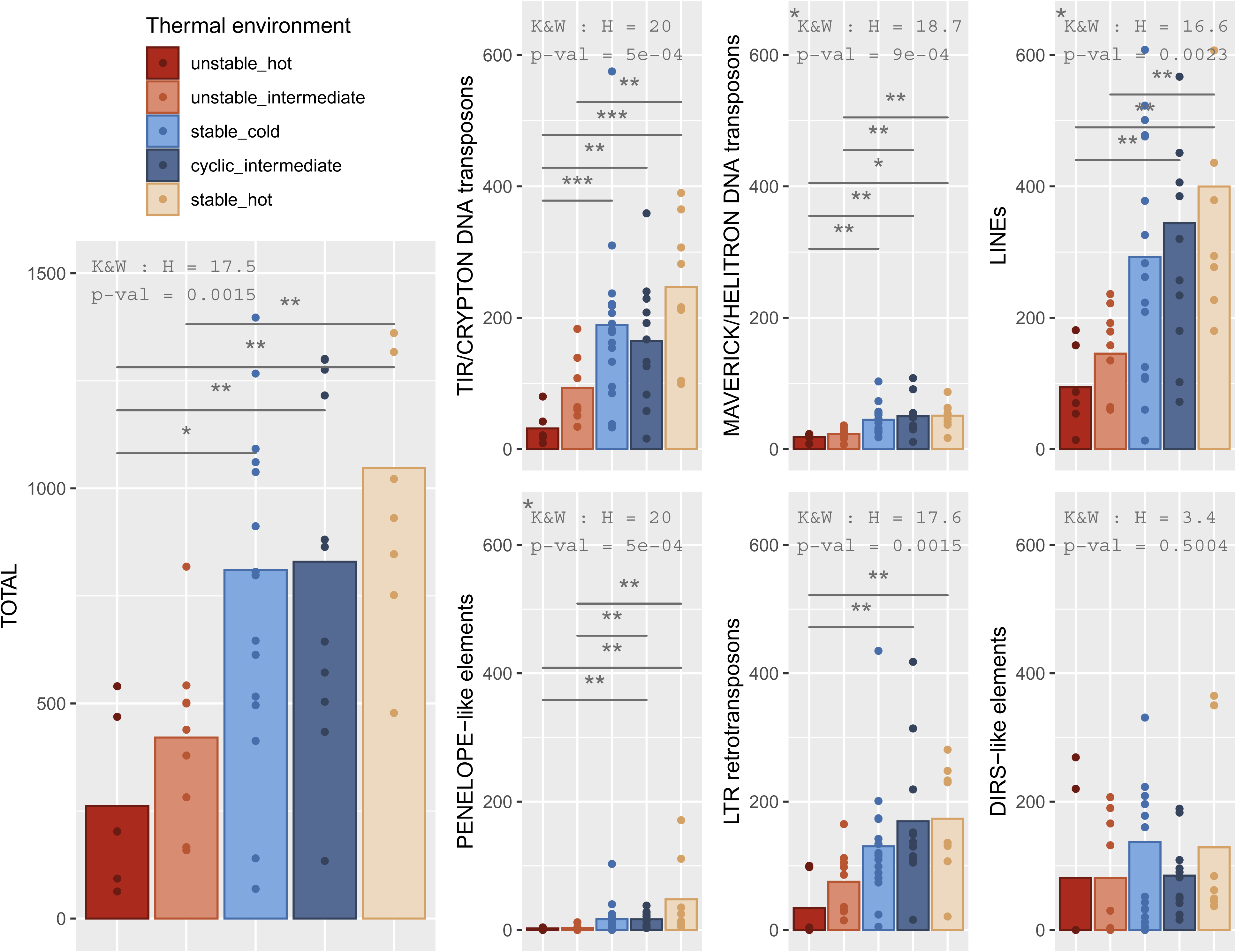
Bar plot of TDF for the six TE orders grouped by thermal environment. For each TE order, the p-value of a Kruskal Wallis test on the effect of the thermal environment on TE diversity is indicated. The stars correspond to the significance level of adjusted result of Nemenyi pairwise tests between every environment (p-values adjusted for multiple tests according to Holm’s correction). An asterisk in the upper left corner indicates the Kruskal&Wallis test was performed on the log transformation of the data to ensure the homogeneity of variance between groups.

### Relationship Between Thermal Environment and Order-Specific TE Diversity

Because each TE order showed drastically different TDF and distribution in polychaetes (Figure 2), we studied the association between TDF and thermal environment separately for each TE order. Except for DIRS-like elements which displayed no significant association, the TDF of all TE orders displayed significant differences between some environmental categories and showed a similar gradient (Figure 4). This consistent trend was statistically supported by the significant conservation of environment’s ranks of TDF across all TE orders (Kendall’s W = 0.88, p-value= 3e-4). The observed gradient of TDF according to the environment was determined primarily by temperature stability and secondarily by mean temperature. Here, the cyclic-intermediate environment was not significantly different from stable environments. Interestingly, TDF in the unstable-hot environment was lower than in the unstable-intermediate environment, while the stable-hot environment displayed higher TDF than the stable-cold or cyclic-intermediate environments. For the five TE orders significantly associated with the thermal categories, the unstable-hot environment which exhibits the lowest TDF, was significantly different from cyclic-intermediate and stable-hot environments, as well as from the stable-cold environment for DNA transposons. However, the lower average TDF of the unstable-hot compared to the unstable-intermediate environment was never statistically supported. For species from the unstable-intermediate environment, TDF was significantly lower than for species in the stable-hot environment, except for LTR-retrotransposons. It also appeared lower than in the stable-cold and cyclic-intermediate environments for each TE order, although these results were not significant for every TE orders. Stable and cyclic environments were not significantly different from one another but showed the same trend across the five TE orders. The stable-hot environment always had the highest TDF, and the cyclic-intermediate environment appeared to be the second regarding TDF, except for TIR/Crypton elements.

To improve our understanding of TDF patterns across thermal environments, we also performed, for each TE order, pGLS regressions that account for the phylogenetic structure of the dataset. Only DIRS-like retrotransposons and TIR/Crypton DNA transposons (Figure 5) showed better AIC with the pGLS model than with the GLS model (Supplementary material 7), although in every case, except DIRS-like, the two AIC where less than 3 units apart, indicating similar performances of the model. The association between thermal environments and TDF was confirmed for Maverick/Helitron DNA transposons, LINEs, Penelope-like retrotransposons, and LTR-retrotransposons (Supplementary material 7). Interestingly, for DIRS, the means of each thermal environments adjusted by the pGLS showed that TDF was clearly lower in unstable than in stable environments, and a significant difference between unstable-intermediate and cyclic-intermediate environments was observed (Figure 5). Moreover, although the means of TDF for TIR/crypton DNA transposons corrected using the phylogenetic signal displayed the same pattern than on Figure 4, differences between environments were close but above the 0.05 rejection threshold (Figure 5). However, the effect of thermal environment became significant after removing the clear outlier *P. stalagmia* (pGLS : R² = 0.45, AIC = 573, p-value = 0.003).

**Figure 5:**
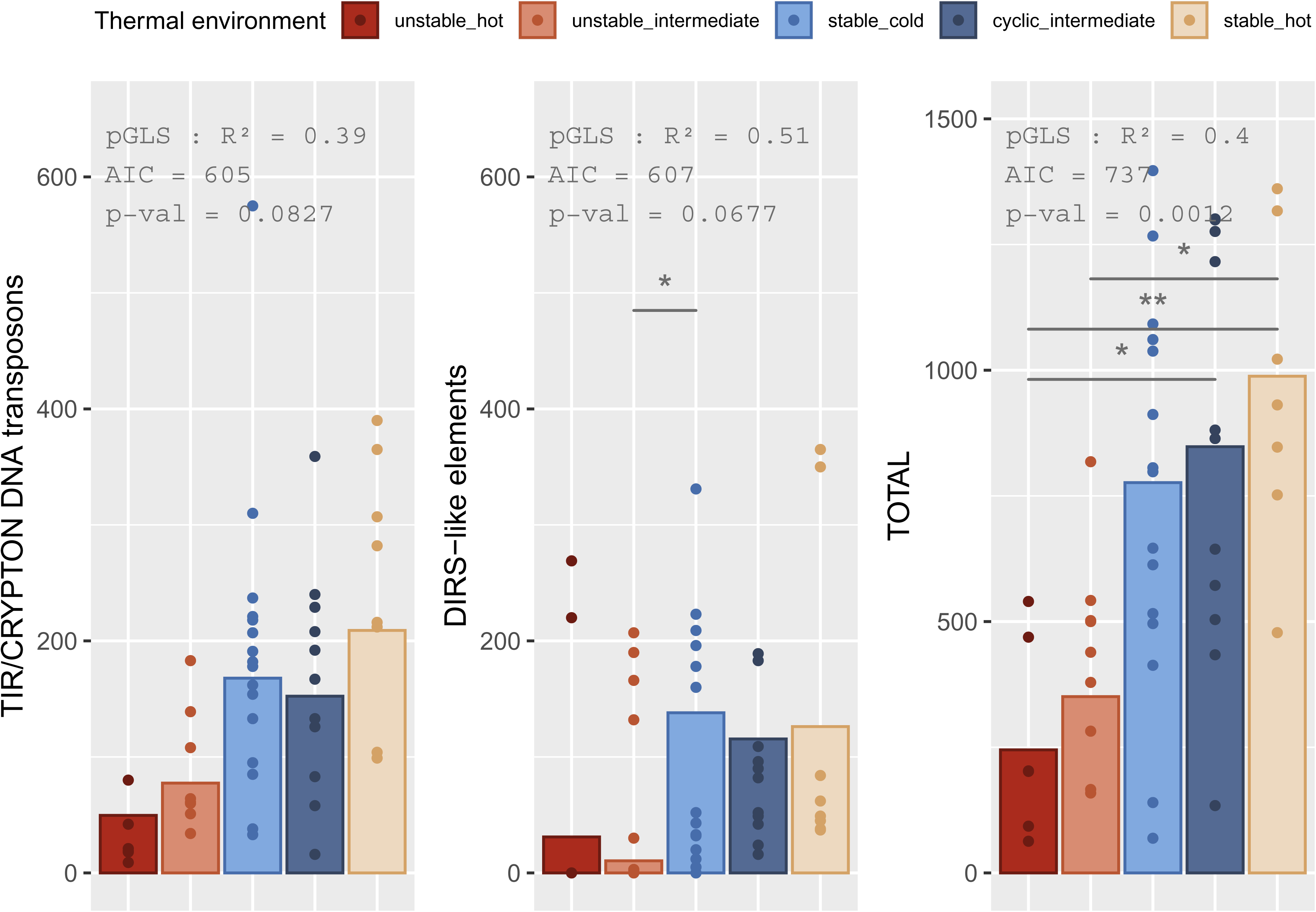
Bar plot TIR/Crypton DNA transposons and DIRS-like elements TDF corrected by pGLS by thermal environments. For each TE order, the R² coefficient, the AIC, and the p-value of the ANOVA on the pGLS are indicated. The stars correspond to the significance level of adjusted result of Tukey HSD tests.

### Exploring the Pattern of DIRS-like Elements TE Diversity

DIRS-like elements showed singular results compared to the other TE orders, both concerning their contrasting TDF between Errantia and Sedentaria species, and their pattern of TDF according to the thermal environment. Thus, we analyzed the two subclasses of polychaetes separately. The diversity of DIRS-like elements in Errantia did not show any relationship with the environment (Figure 6). In contrast, a significant relationship between TDF and the environment was found for Sedentaria. Interestingly, in this case, the same trend described above for the other TE orders was again observed. DIRS-like elements of Sedentaria showed lower diversity in unstable environments associated with hydrothermal habitats. Notably, no DIRS-like elements were detected in any of the four Sedentaria species sampled from the unstable-hot environment. Another particularity is that species from the cyclic-intermediate environment displayed the highest DIRS-like family numbers, although these were not significantly different from those in stable environments.

**Figure 6:**
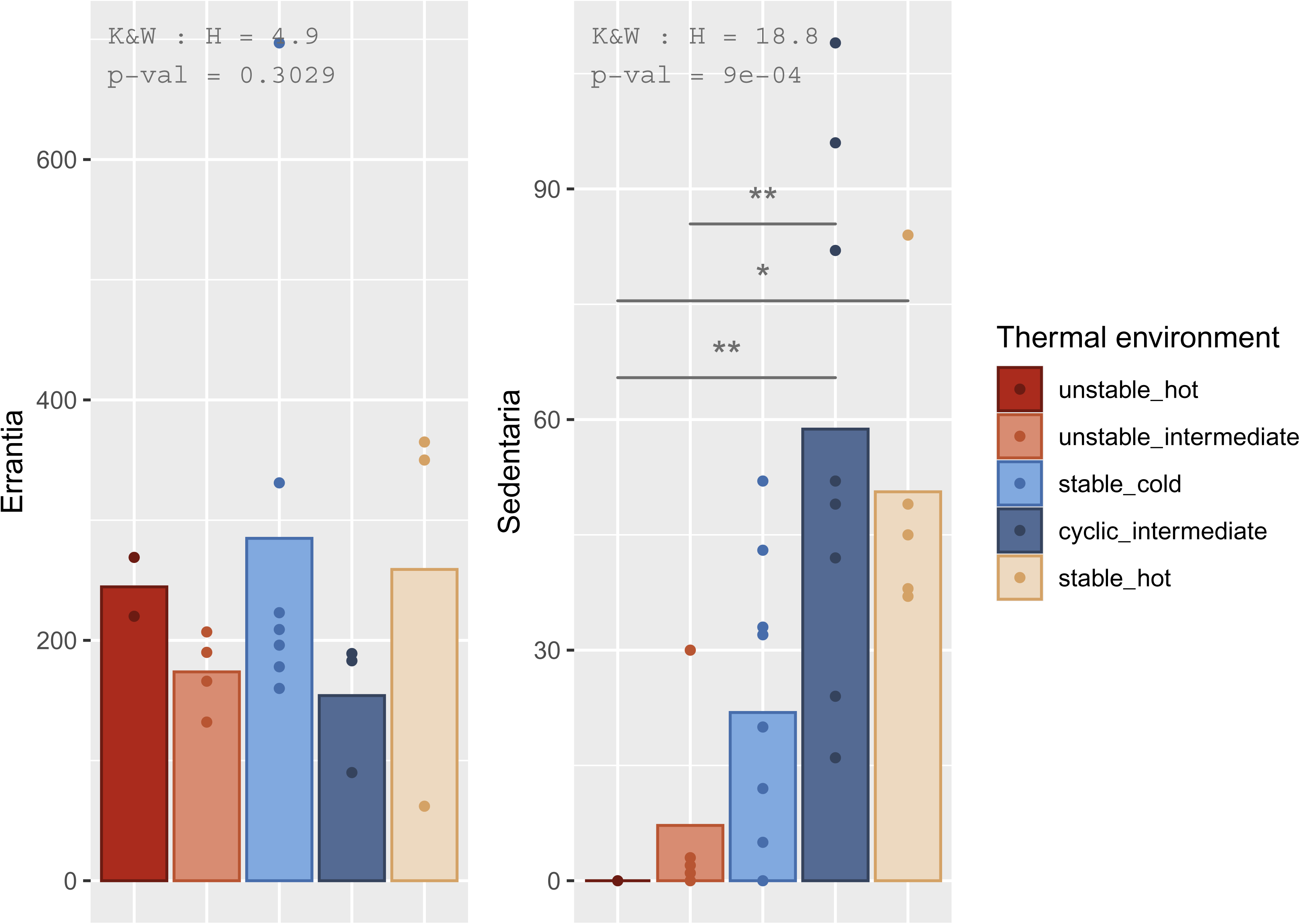
Bar plot of DIRS-like diversity, grouped by thermal environment with A) only Errantia species and B) only Sedentaria species. For each TE order, the p-value of a Kruskal Wallis test on the effect of the thermal environment on TE diversity is indicated.

## DISCUSSION

### DetecTE, an Optimized TE Detection Tool

To date, very few studies have focused on the relationships, between the TE diversity of family and environmental characteristics of host’s habitats for marine animals, on the basis of a large dataset of species. In this framework, the present work is a pioneering study for polychaetes with such a large sample covering a deep phylogeny of these organisms. Because complete genomes are scarce in this phylogenetic group, we used a transcriptomic approach whose efficiency and reliability to assess TDF have already been demonstrated (Filée et al,2021). In addition, we took advantage of recently available whole genomes of the polychaete species in our dataset to compare TDF estimates from whole genomes and transcriptomes from the same species. The higher TDF we observed in genomic data compared to transcriptomic data was expected, since inactive TE copies obviously escape transcriptomic surveys. However, our purpose was to investigate the potential link between TDF and environmental conditions. We thus operated within the paradigm that the higher the TDF is in the genome, the higher it is in the transcriptome, as also supported by our data (Supplementary material 5), and that TDF is in turn modulated by natural selection and contributes to the whole genome mutagenicity (see further sections of the discussion). In that sense, families whose copies are no longer autonomous or only very weakly expressed are therefore not the focus of this paper.

We chose to update our previous tool (DetecTE1) for TE annotation in transcriptomes without reference genomes. This first version of this tool was suitable for characterizing TE families in polychaetes (Filée et al., 2021) but was assumed to overestimate the number of TE families. This overestimation was most likely because copies from a given family can be assembled into multiple transcripts, in addition to isoforms, which can lead to them being counted as separate families. To mitigate this bias, DetecTE2 allows users to merge sequences belonging to the same family based on their similarity. In DetecTE2, TE sequences belonging to different superfamilies but grouped on the same transcript are however separated, allowing a more precise and complete estimation of TDF. Furthermore, DetecTE2 can process data from multiple assemblers, tissues, or individuals simultaneously to maximize detection. Overall, although only a few polychaete whole genomes are currently available, we demonstrate that a transcriptomic approach is a powerful, reliable, and feasible method for estimating the diversity of active TE families. This approach is particularly well-suited for studies on a wide range of non-model species, given the currently available data, or species with large genomes, such as krill, for which whole genomes are still challenging to sequence and assemble today because of their large sizes (Jeffery, 2012). We believe that DetecTE2 is an essential tool to advance the study of TEs across the tree of life.

### The Influence of Polychaete Phylogeny on TDF

Since TEs are essentially vertically transmitted (Wallau et al., 2016), we can reasonably expect the host phylogeny to have a significant impact on TE distribution and TDF, and it has indeed been described in many contexts. For example, retrotransposons appear to be restricted to eukaryotes (Boeke, 2003), and Maverick elements have never been detected in plants (Pritham et al., 2007). At the superfamily level, TE diversity can vary depending on the taxa. For instance, Copia LTR-retrotransposons display low diversity in metazoans but are highly diverse in plants.

For each TE order, to consider the potential influence of phylogeny on TDF, we used pGLS, which produced better fits to the data than regular GLS for only TIR/Crypton DNA transposons and DIRS-like elements. This result is not surprising for DIRS-like elements, considering the significant difference of TDF between Errantia and Sedentaria for this order. In particular, Errantia exhibited higher TDF for both DIRS and Ngaro elements (Supplementary material 6), the two superfamilies of the DIRS-like order. This suggests that if a strong increase in TE diversity, and most likely in copy numbers, occurred in Errantia, it has probably affected most of the tyrosine recombinase retrotransposons, that are characteristic of the DIRS-like order. This is particularly interesting considering that these two superfamilies of elements have quite different structures, such as the presence or absence of a methyltransferase and the nature of the sequences involved in transposition (Poulter & Butler, 2015). The DIRS-like TDF in Sedentaria is consistent with that of animals in general (0 to 81 families; Piednoël et al. 2011), suggesting a specific increase in Errantia rather than a decrease in Sedentaria. However, few studies on the distribution and dynamics of DIRS-like elements exist, preventing further hypotheses on the mechanisms underlying this difference between the two subclasses.

For TIR/Crypton order, we observed lower TDF in Alvinellidae compared to Terrebellidae, both belonging to Terrabellida order. Similarly, Lepidonotopodinae species exhibited lower TDF compared to other species of the Phyllodocida order, which explains the improvement of the fit when using a pGLS. Nonetheless, in both cases, there was a correlation between the thermal environment and the tree topology. In our dataset, all Alvinellidae species, and all Lepidonotopodinae species except *B. jasoni,* are indeed from unstable environments, while species in Terrebellidae family and the rest of the Phyllodocida species are from stable environments. In fact, the structure of our species sample did not allow us to fully disentangle the respective contributions of the environment phylogenetic lineages history to TDF, due to the inherent non-zero covariance between these two factors. A more precise correction of TDF using pGLS could be obtained with a larger dataset, or by including the actual phylogeny of our dataset in the model. It is, however, currently unavailable and challenging to generate, due to the presence of closely related species and lineages that diverged 500 Mya in the same dataset (H. Chen et al., 2020).

For other orders, considering phylogenetic relationships between species did not improve the regression model fit. This, however, does not exclude the possibility that the evolutionary history of species may have influenced their TDF at lower levels of TE classification. For example, the GalEA clade of TE (which comprises closely related elements of the Copia superfamily), or the Suzu and Simbad clades (Bel superfamily), are indeed restricted to Sedentaria species (Filée et al., 2021). Differences at these lower levels may be masked when studying TE orders or overall TDF.

At a much lower level of polychaete phylogeny, the case of the genus *Osedax* appears singular. The two *Osedax* species were clearly distinguishable from other species from the stable-cold environment, exhibiting the lowest TDF by far. Notably, the next species from the stable-cold environment with the lowest TDF is *Microspio moorei*, which has almost three times as many families. Even though only two *Osedax* species were studied, their similar and unusually low TDF may indicate a property specific to this genus. Indeed, *Osedax* species, also called “bone-eating worms,” live exclusively on sunken marine mammal carcasses (Rouse et al., 2004; Vrijenhoek et al., 2009), and use obligate bacterial symbionts to digest the bones and nourish themselves (S. K. Goffredi et al., 2005). Most likely as a consequence of its adaptation to this highly specialized lifestyle, *Osedax* species have undergone drastic genome size reduction affecting both genes and transposable elements (Moggioli et al., 2023). Some metabolic functions were lost and are now provided by their bacterial symbionts, which notably harbor a higher percentage of TEs than their free-living relatives (S. Goffredi et al., 2022). In fact, *O. frankpressi* has the smallest genome among the 23 species studied with known genome sizes (Supplementary Material 1). The particularly low TDF in *O. frankpressi* and *O. japonicus* is probably linked to this genome reduction, which appears independent of temperature in the deep sea. Regardless, with or without these two species in the dataset, all statistical tests showed similar results, highlighting the robustness of the present observations.

### The link between genome size and TE diversity

If genome size is known to be positively correlated with the absolute abundance of TEs in eukaryotes (Chénais et al., 2012; Elliott & Gregory, 2015; Gregory, 2005; Marino et al., 2024), the links between genome size and TE diversity are not yet fully characterized. Most theoretical models considering the relation between the 1C genome size value and the diversity of TE, predict a negative correlation (for a summary, see Wang et al. 2021) because of selection against ectopic recombination and competition between TE families in large genomes (Boissinot & Sookdeo, 2016; Haley & Mueller, 2022; Petrov et al., 2003). However, empirical studies of genome size and TE diversity correlations yield contrasting results. Indeed, positive, negative or even no correlations have been reported depending on the lineage studied and the chosen phylogenetic scale (Elliott & Gregory, 2015; Petersen et al., 2019; Zuo et al., 2023; Decena-Segarra & Rovito, 2024), which prevent us from concluding. To clarify this in the annelid species in the present article, and add to the knowledge on the potential influence of TE diversity on genome size, we computed the correlation on the basis of a subsample of 23 species whose genome size has already been estimated (Filée et al. 2021, www.genomesize.com). No significant correlation was observed (spearman correlation: ρ = 0.341, S = 1332.8, p-value = 0.111; Supplementary material 8). Thus, the observed pattern of TDF across habitats and across the phylogeny of polychaetes does not affect polychaetes genome size in principle.

### The influence of the thermal environment on the TDF

The results reported in this paper showed a clear relationship between the transposable elements diversity of family and environmental conditions in which the annelid species live. The diversity of TEs within genomes is not only the product of progressive genetic divergence through time. TE diversification is, in fact, a very complex and still not a fully elucidated process (Wells & Feschotte, 2020; Pulido & Casacuberta, 2023), resulting from the association of several mechanisms acting at the genome and the population level.

TE evolution, and thus diversity, is driven by evolutionary forces such as drift, itself impacted by population demography, gene flow, and selection (Lockton et al., 2008; Brunet & Doolittle, 2015; Szitenberg et al., 2016, discussed below). Furthermore, within genomes, TDF can increase by the insertion of new copies, which may diversify rapidly through the accumulation of mutations or the acquisition of new functional domains, and give rise to new TE families (H. Eickbush & K. Jamburuthugoda, 2008; Lerat et al., 1999; Wells & Feschotte, 2020). Element diversity may also increase through inter-element recombination between genomics copies, above all during transposition of LTR-retrotransposons by reverse-transcription-related recombination (see Drost & H Sanchez, 2019 for review). Inter-element recombination could also be a shared property of diverse TEs, as it has been observed for specific non-LTR retrotransposons, like SINEs elements, in protist (Yadav et al., 2012). Similarly, DNA transposon chimerization and diversification was shown in lizards or *Daphnia* (Novick et al., 2011; Vergilino et al., 2013). Plus, as a result of successful transposition bursts in *Saccharomyces cerevisiae*, chromosomal copies of newly inserted element revealed contributions from both young and older family members, capable of generation of TE copies that are a mosaic of TE from different families. However, several molecular mechanisms, such as ectopic recombination between TE copies, and subsequent deletions (Ji & DeWoody, 2016), TE silencing by the host’ genome (Sammarco et al., 2022) and copy inactivation by mutation (Blumenstiel, 2019), are key factors counteracting the expansion of TE copy numbers and limiting TE diversity. Also, competition between diverse TE families for the host’s replication machinery (Furano et al., 2004) or opportunities to escape silencing processes contribute to the evolution of TE diversity at the genomic level (Abrusán & Krambeck, 2006). Overall, these mechanisms may lead to the extinction of some TE families at the genome scale and therefore the reduction of TDF.

In addition to the diversification or decay of existing TE families within genomes, horizontal transfers of transposable elements (HTT) can bring new families to the genome and consequently increase TE diversity. HTT have major consequences on the evolution of eukaryotes (Schaack et al., 2010). But, if they play an important role in maintaining TEs in eukaryotic genomes (Venner et al., 2017), the diversity of TE families within a species derives mainly from the evolution of genome copies and only to a minor extent from HTT (Muller et al., 2025). In the case of polychaetes in the present study, more than 95% of the detected TE families were species specific (data not shown). It is therefore likely that HTT has only marginally contributed to the variation of TDF among species.

Interestingly, when comparing TDF between environments, all significant differences were observed between thermally unstable environments on one hand, and cyclic or stable environments on the other hand. Besides, the observation of a similar pattern in all TE orders (except for DIRS-like elements in Errantia), despite their drastically different structures and dynamics, suggests the existence of a mechanism affecting TEs as a whole and revealing variations between habitats. Unstable environments in this study correspond to hydrothermal vents, which are also peculiar environments for many reasons. It is therefore challenging to identify the factors that can explain the observed pattern of TDF. However, a notable aspect of hydrothermal vents in comparison with other habitats is that they are highly dynamic habitats, submitted to recurrent extinctions and volcanic events (Vrijenhoek, 2010). Hydrothermal species, at least in their adult forms, display a patchy distribution and experience repeated loss of habitats and local demographic bottlenecks due to new colonization events , thereby reinforcing genetic drift effects and possibly reducing genetic diversity (Coykendall et al., 2011; Maruyama & Kimura, 1980). Although the extinctions are local and only affect a small proportion of the species’ general population, their effect may affect the whole species. In particular, it can contribute to the existence of genetic differentiation between populations (Sgarlata et al., 2025), as already shown in some species (Jang et al., 2016; Xi et al., 2023).In these conditions, TDF may vary between hydrothermal populations and the estimated divergence between species for TDF would then depend on the design of sampling. However, this scenario can hardly explain the pattern of interspecific TDF we observed. It is difficult to conceive that neutral evolutionary mechanisms alone would result in significantly lower TDF values in the 13 hydrothermal species compared to those sampled in other habitats, especially since the same pattern is observed across different TE orders. Additionally, because phylogenetic constraints contribute to TDF only in TYR/Crypton elements, and given that hydrothermal species are widely distributed across the phylogenetic tree of our sampled species, a common ancestral heritage of lower TDF in hydrothermal species compared to species in other habitats is unlikely. Nevertheless, in the absence of robust inferences regarding the population demographic histories and population genetic structure of the studied species, we cannot exclude the possibility that more complex patterns of species demographic history might align with our results. Indeed, pioneering studies on this issue for hydrothermal species have revealed stark contrasts in the level of genetic diversity and in population genetic structure between the studied species (Breusing et al., 2023; Nakajima et al., 2024).

We instead propose that selection could be the main factor explaining the difference of TDF across environments. In that case, the lower number of TE families in hydrothermal species might be explained by one of the many physical and chemical characteristics of hydrothermal vents, such as the high pressure associated with depth or the high concentration of reduced sulfur compounds and other toxic chemicals linked to chemosynthesis-based ecosystems (Campbell, 2006; Canganella, 2001). However, abyssal species that are also subject to high pressure, or species from other chemosynthetic habitats such as cold seeps (e.g. *Lamellibrachia satsuma*, *P*. *echinospica*, and *Harmothoe sp*.) where gas emerges from sediments (Ceramicola et al., 2018), have clearly more TE families than hydrothermal species. Furthermore, other environmental factors might cause, alone or in conjunction with others, this difference of TDF between hydrothermal vents and other habitats. Among them, we argue temperature is a good candidate for different reasons. First, our results showed the existence of a consistent pattern of TDF variation across TE orders, correlated to the thermal properties of the habitats occupied by annelid species. Furthermore, although the species from the two hydrothermal environments, hot and intermediate, do not differ statistically in their level of TDF, they are distinguished from each other notably in their contrasts with other environments. Second, the effect of temperature on TE activity and transposition has been largely studied (Capy et al., 2000; de la Vega et al., 2007; Guerreiro, 2012; Jardim et al., 2015; Rose et al., 2016; J. E. Chen et al., 2018; Bogaerts-Márquez et al., 2021). One way for TEs to transpose is through the acquisition of regulatory sequences whose function is to recruit transcription factors that are sensitive to stress conditions (Vernhettes et al., 1998; Hermant & Torres-Padilla, 2021; Mackey et al., 2025). Thus, the composition of the genome in TEs appears to result from an adaptative response to different host-specific stimuli and might reflect the environmental history of each host (Grandbastien et al., 2005). The ONSEN retrotransposon (Cavrak et al., 2014) or the Tam3 DNA transposon (Hashida et al., 2006) are two examples of this feature in the case of heat stress. Because of stress-sensitivity, repeated temperature changes can lead to bursts of transposition (Belyayev, 2014; Castanera et al., 2020; L. Chen et al., 2019) which can increase TDF *via* the diversification of copies over time. In return, with increased TE diversity, the possibility that some elements will escape regulatory systems also increases. It is therefore hypothesized that the transposition rate increases with the TDF in a genome. However, if some mutations due to new TE insertions may be beneficial and can contribute to evolutionary innovations, most of them are expected to be neutral and, above all, deleterious (Barrón et al., 2014; Arkhipova, 2018). In other words, high TDF, by enhancing transposition, might promote some adaptative mutations while increasing the mutation load, that may end with population extinction.

Thus, our hypothesis is that a hypervariable habitat such as hydrothermal environments may be highly mutagenic because of transposition activation in response to stress. In such habitats, high TDF might be counter selected because of the accumulation of mutations, a large proportion of which being deleterious (Figure 7). Conversely, in stable environments that do not often trigger TE transposition, a TDF that is too low could be counter-selected because of the low ability to produce adaptative mutations. We therefore argue that the population’s mean TDF values, itself impacting the transposition rate, can be seen as the result of stabilizing selection on the mutation rate. This selective pressure would lead to different TDF, depending on the variability of the environment, because of stress-sensitivity of TEs and the link between TDF and transposition rate. This process would allow populations to maintain a level of TDF ensuring compatibility with both population survival and the maintenance of population adaptive potential.

**Figure 7:**
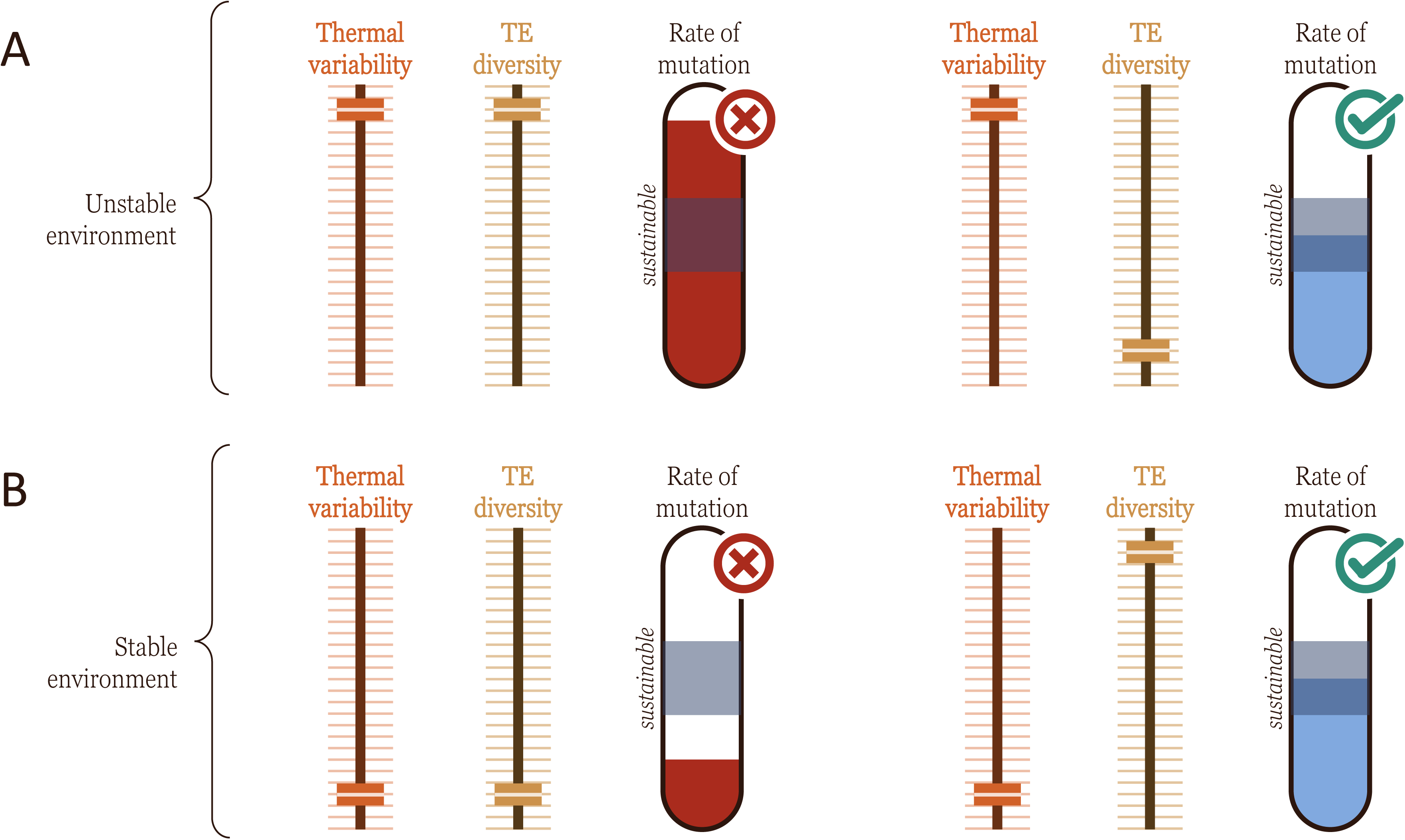
Graphical representation of the selection hypothesis. Rate of mutation depending on the TE diversity in A) unstable and B) stable environments. The reported mutation rates are not quantitative. Thermal variability and TE diversity are assumed to be positively correlated with the mutation rate, which is under selection to remain in a “sustainable” range at the population level.

An alternative hypothesis, which is not mutually exclusive with preceding one, is that competition between TE families leads to a decrease in TE diversity in case of strong mobilization events of some TE families. Theoretical models predict that in presence of cross-reactive regulation mechanisms, such as siRNA systems (Rigal & Mathieu, 2011), the high mobilization of some TE families, because of a stress for example, induces a higher global silencing of TEs, which decreases the activity of all other families and lead to a decrease in TE diversity (Abrusán & Krambeck, 2006). Consequently, the TDF would be lower in unstable environments, that strongly trigger the stress sensitive families, compared to cyclic and stable ones.

According to our proposal, the optimal TDF value is affected by processes operating at the genome level, creating competition between TE families and limiting their diversity, and then at the population level, by selecting individuals whose TDF correspond to the best fitness. These mechanisms shape TDF differently depending on thermal stability, and more specifically on the frequency of exposure to stress.

## Conclusion

This first large-scale study of TE diversity (estimated at the TE family level), in relation to the thermal conditions of life, with species from the same taxonomic group, the polychaetes, revealed significant variability across contrasted thermal environments. These differences mainly suggested a strong influence of the thermal stability parameter rather than that of the average temperature. To confirm these findings and test the evolutionary model we proposed, it would be valuable to explore new taxa to determine whether this pattern is taxa-specific or can be generalized, at least in marine animals. The major constraint remains the need for sufficient omics data to clearly represent several contrasted environments, while including phylogenetically close species. Decapoda would be another suitable model, as they exhibit numerous species in all the environments we studied here, with a large amount of transcriptomic data already published. In complement to the TDF, studying the influence of thermal environment on TEs percent in genomes, or TE copy numbers, in marine species, would be useful to improve our understanding of the process by which species may regulate their global TE-mediated mutation load. However, this would require a large number of assembled genomes or low-coverage sequencing.

## Acknowledgment

The authors would like to thank Sébastien Duperron and Éric Thiébaut for their valuable help with diversity analyses and statistics. We also thank the ABIMS bioinformatics platform for providing the necessary software and computational infrastructure for the analyses. Special thanks go to Charlotte Berthelier and Erwan Corre for their advice on the management of transcriptomic data and good coding practices. We would also like to extend our thanks to Didier Jollivet, François Lallier, and Gregory Rouse for their valuable help in assigning polychaete species to thermal environments. Finally, we sincerely thank Jonathan Filée and Denis Roze for their help in reviewing the manuscript.

## Data availability

No sequencing data were generated for this article. References for accessing the data are provided in Supplementary Material 1.

## Benefits Generated

Benefits from this research accrue from the sharing of our results publicly.

## Authors contribution

E.B. and L.L. designed the research ; S.H. and L.L. constructed the dataset and established metadata ; L.L. processed the transcriptomes and developed the pipeline ; L.L. and T.R. performed statistical analyzes ; E.B., L.L. and T.R. wrote the manuscript ; E.B., S.H., L.L. and T.R. validated the manuscript ; E.B. supervised the project.

