## Supplementary figures and images for "Taming the Genetic Fire: Transposable Element diversity across thermal environments in polychaetes"

### Supplementary material 3

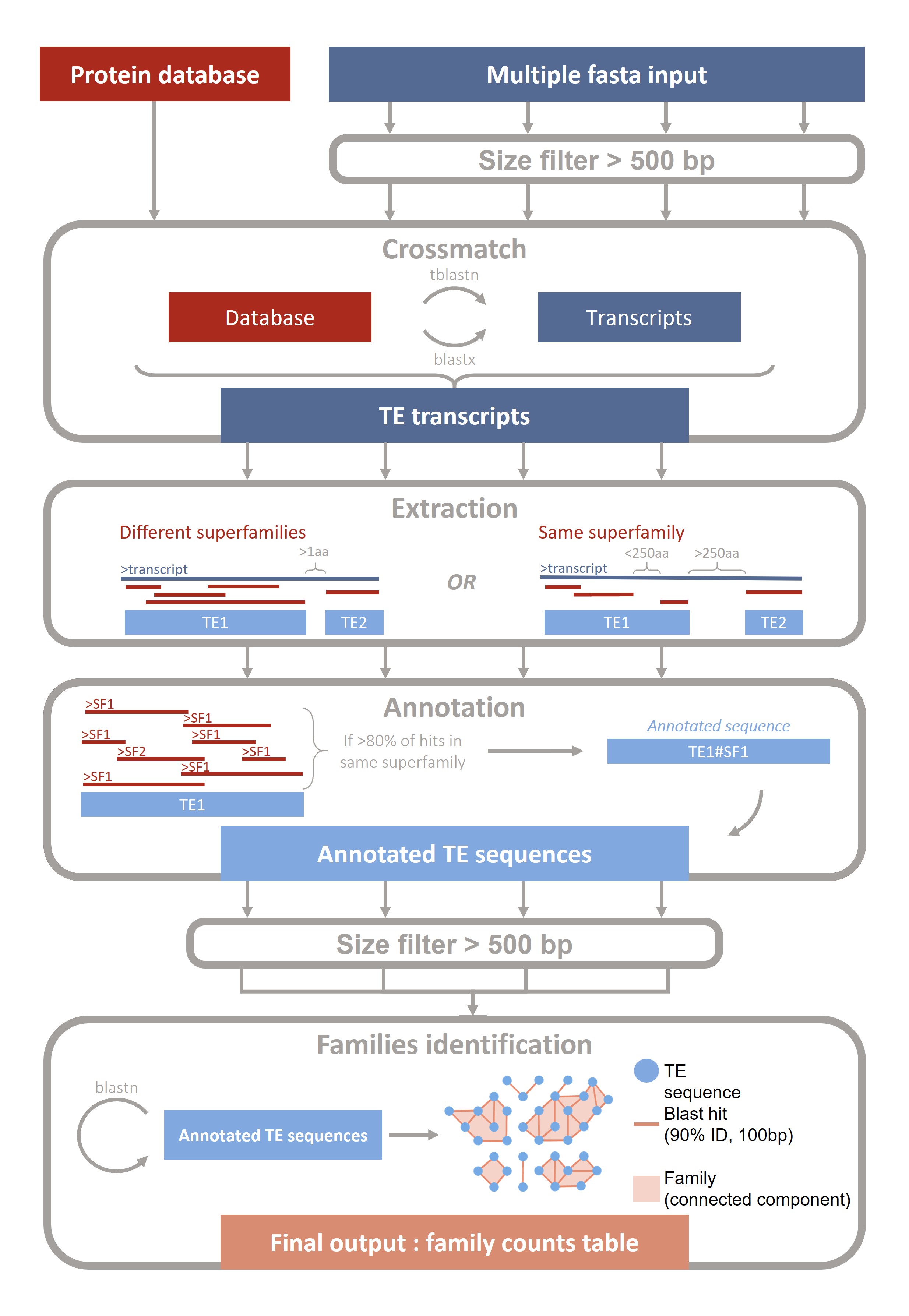

### Supplementary material 4

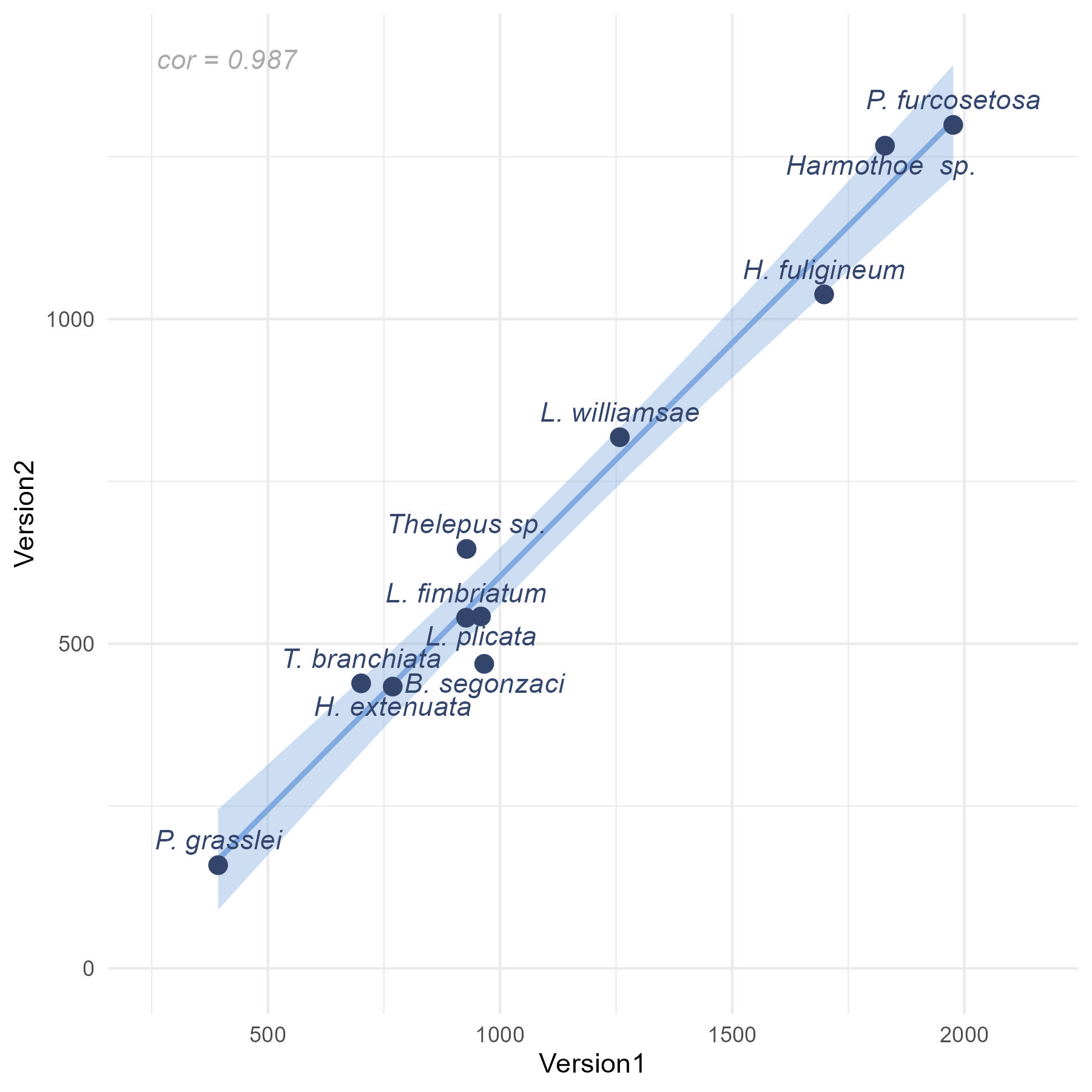

### Supplementary material 8

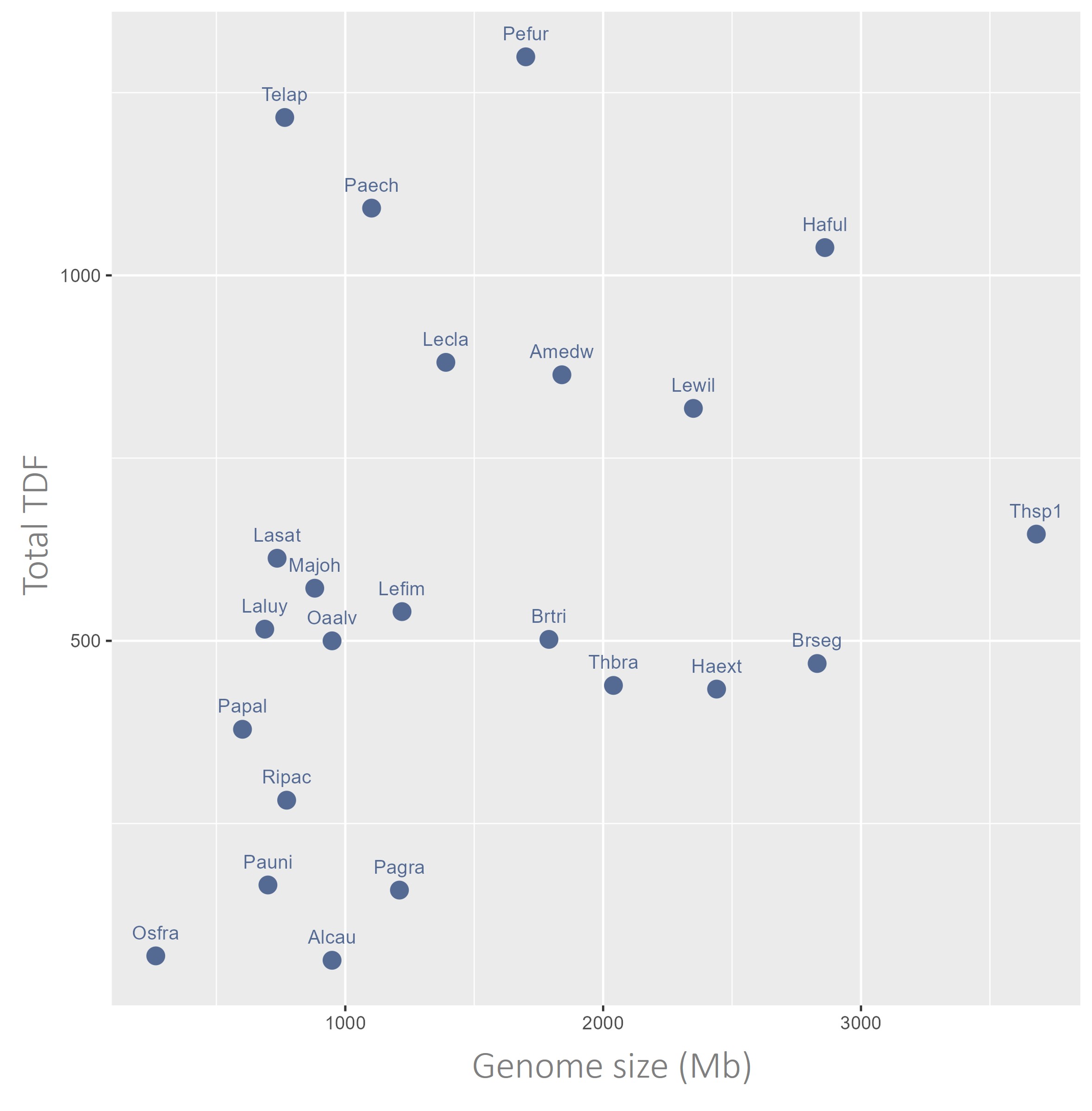
